# Insulin stimulated upregulation of OCTN2 carnitine transporters is impaired in patients with Primary carnitine deficiency

**DOI:** 10.1101/2025.09.22.677318

**Authors:** Rannvá Dahl, Tórur Sjúrðarson, Poula Paturson, Arthur Ingersen, Peter F. Andersen, Jonas E. M. Jørgensen, Elizabeth J. Simpson, Jan Rasmussen, Kasper Kyhl, Paul L. Greenhaff, Flemming Dela, Steen Larsen, Clara Prats

**Affiliations:** Xlab, Center for Healthy Aging, Department of Biomedical Sciences, Faculty of Health Sciences, University of Copenhagen, Copenhagen, Denmark; Department of Internal Medicine, The National Hospital of the Faroe Islands, Tórshavn, Faroe Islands; MRC-ARUK Centre for Musculoskeletal Ageing Research, ARUK Centre for Sport, Exercise, and Osteoarthritis, School of Life Sciences, The Medical School, University of Nottingham, Nottingham, United Kingdom; Clinical Research Centre, Medical University of Bialystok, Bialystok, Poland; Department of Cardiology, Rigshospitalet, Copenhagen University Hospital, Denmark

## Abstract

**Background:** Primary carnitine deficiency (PCD) is an autosomal recessive disorder characterized by a lack of functional carnitine transporters OCTN2 (Organic Cation/Carnitine Transporter 2), which has been linked to several cases of sudden death in young Faroese individuals. It causes low carnitine levels and can present with hypoketotic hypoglycemia, skeletal and cardiac myopathy. Patients are treated with L-carnitine, and even while receiving treatment, skeletal muscle carnitine levels are only approximately 7% of normal. The regulation of the carnitine transporter, OCTN2, is not fully understood, but a combination of hypercarnitinaemia with hyperinsulinemia upregulates skeletal muscle carnitine uptake and OCTN2 mRNA expression.

**Objective:** The question arises as to whether hyperinsulinemia increases the efficiency of L-carnitine supplementation in PCD. The present study investigates the regulatory mechanisms behind insulin-induced carnitine uptake, and whether the combination of hypercarnitinaemia and hyperinsulinaemia increases skeletal muscle carnitine levels in patients with PCD. In addition to our main goals, we also explored the measurement of whole-body fat oxidation at rest and during exercise

**Method:** Nine patients with PCD (homozygous for the c.95 A > G, pN32S mutation) and nine healthy controls matched to age and body mass index (BMI) participated in the study. A six hour hyperinsulinemic clamp was supplemented with infusion of L-carnitine the last five hours. Skeletal muscle biopsies were collected before and after the clamp and carnitine content was measured. Furthermore, confocal microscopy was used to access regulation of OCTN2 and GLUT4 positive vesicles due to insulin stimulation and additionally whole body fat oxidation was measured with indirect calorimetry.

**Results:** We found that the combination of hypercarnitinaemia with hyperinsulinemia did not increase skeletal muscle total carnitine levels significantly for neither patients with PCD [4.5 (SE 0.6) to 5.3 (SE 0.5) mmol · kg^-1^] (*P* = 0.28) nor controls [19.8 (SE 0.6) to 21.2 (SE 0.5) mmol · kg^-1^] (*P* = 0.053). The muscle carnitine profile showed that patients with PCD have low levels of total and free carnitine in skeletal muscle, but normal levels of acetylcarnitines corresponding to 60% of total carnitine (normal is ∼16% of total muscle carnitine). The results from confocal microscopy indicate that insulin regulates skeletal muscle carnitine uptake by stimulating OCTN2 recruitment from intracellular storages to the plasma membrane. This regulatory mechanism is however impaired in patients with PCD. Furthermore, we found that PCD patients were more dependent on carbohydrates at rest. In regard to fat oxidation, no difference was found between PCD and control group during short-term exercise.

**Conclusions:** The study indicates that insulin stimulates translocation of OCTN2 to the plasma membrane in healthy controls, a mechanism that seems to be impaired in patients with PCD. The combination of hypercarnitinaemia with hyperinsulinemia did not increase skeletal muscle total carnitine levels significantly (*P* = 0.28) in patients with PCD.

## Introduction

For several decades, multiple cases of sudden death in young Faroese individuals with untreated primary carnitine transporter deficiency (PCD) have been reported [1-3]. The prevalence of undiagnosed PCD remained unknown until 2009, when a nationwide voluntary screening program was initiated and all Faroese inhabitants were invited to have their blood carnitine levels analyzed [4, 5]. The screening indicated that the prevalence of PCD in the Faroe Islands is the highest reported worldwide with a prevalence of 1:300 [4, 6], compared with approximately 1:40,000 – 1:100,000 worldwide [7, 8]. PCD is an autosomal recessive disorder that is characterized by mutations in carnitine transporters (i.e., the organic cation transporter 2 (OCTN2)), reducing the level of stored carnitine. Untreated PCD has been connected to several incidences of sudden cardiac death of young Faroese individuals, but the underlying pathogenic, cellular mechanisms in patients with PCD remained elusive.

More than 95% of the body’s total carnitine store exists within skeletal muscle cells as either free or acylcarnitines which are esters arising from the conjugation of activated long-chain fatty acids (fatty acyl-CoA) with free carnitine [9, 10]. Fatty acyl-CoA groups can not directly cross the mitochondrial inner membrane to enter the mitochondria and acylcarnitines play an essential role in transporting fatty acyl-CoA groups across the membrane and into mitochondria matrix for β-oxidation, leading to production of energy [11-14]. Over the past three decades, many additional actions and roles of acylcarnitines have been discovered. For instance, it is clear that they are also involved in maintaining the homeostasis of the fatty acetyl-CoA/free CoA ratio at the onset of exercise [15, 16]. Mitochondrial fatty acetyl-CoA is generated from β-oxidation of fatty acids and pyruvate from glycolysis; if the rate of acetyl-CoA generation exceeds the rate at which it is utilized in the citric acid cycle, fatty acid β-oxidation may be inhibited. Free carnitine acts as a buffer by reacting with acetyl-CoA to form acetylcarnitine and thus acetyl-CoA is eliminated [9, 15, 17]. Heart muscle is especially vulnerable to carnitine deficiency since β-oxidation is the primary energy supply [18]. L-carnitine supplementation has shown to be an efficient treatment for PCD [19-21], despite the fact that skeletal muscle carnitine concentration is only 7% compared to healthy controls [22, 23]. Given the low levels of carnitine in skeletal muscle, it raises the question as to how L-carnitine supplementation can have such a positive effect on morbidity and likely mortality in PCD patients.

Carnitine transport regulation is not fully understood, but Stephens et al. showed that hyperinsulinaemia increased total skeletal muscle carnitine content by 13% in healthy controls after hypercarnitinaemia, and was associated with a 2.3-fold increase in OCTN2 mRNA expression [24, 25]. Moreover, both GLUT4 and OCTN2 have highly conserved N-terminal sequences and when their amino acid sequences are aligned, we can observe that they share the critical phenylalanine residue postulated to be essential for insulin stimulation in GLUT4 [26, 27], suggesting that OCTN2 may also be adaptive to insulin. This prompts the question arises of whether insulin regulates skeletal muscle carnitine uptake in a similar fashion as glucose uptake, possibly by stimulating the recruitment of intracellular transporter pools to the plasma membrane [28, 29] and whether such a regulatory mechanism is impaired in PCD or can be used to improve the efficiency of L-carnitine supplementation in PCD treatment. The present study was designed to investigate whether the combination of hypercarnitinaemia and hyperinsulinaemia increases skeletal muscle carnitine uptake in patients with PCD, and whether insulin-induced upregulation of skeletal muscle carnitine uptake is dependent on sarcolemmal recruitment of OCTN2.

## Materials and Methods

### Participants

Eighteen Faroese subjects (18-46 years old) were included in our study. Nine patients with PCD (homozygous for the c.95 A > G, p.N32S mutation), and nine healthy subjects who were matched by sex and age. The number of identified Faroese PCD patients, homozygous for the p.N32S mutation and between 18 and 80 years old, was 28 at the time of the study. They were all invited to participate in the study and nine patients accepted the invitation (Fiure 1). One PCD patient and one control did not complete the study. They were both female and the same age and body mass index (age 22/23 years; BMI 25/23 kg/m^2^). The patients were all taking their daily, individually adjusted, L-carnitine supplementation to keep total free carnitine in blood at a concentration >20 µM, which was monitored with regular blood measurements [22].

### Study approval and registration

All methods and procedures of the present study were approved by the Faroese Ethics Committee and the Data Protection Agency and in accordance with the Declaration of Helsinki. The purpose, nature and potential risks of the study were explained verbally and in writing before individuals gave their informed consent to participate in the study.

### Study design

Participants reported to the laboratory on two different occasions, 5-8 days apart. In both occasions the subjects came overnight fasted (∼10 hours), however L-carnitine supplemented. Moreover, the subjects had abstained from strenuous exercise for 24 h (0). On the first day the participants’ health status was determined by a physical examination which included medical history, an electrocardiogram, a full blood count, electrolyte assessment and PCD (c.95 A > G) genotyping which was followed by a DXA scan and an exercise test to determine fat oxidation, and maximal oxygen uptake (VO_2_peak). On the second day, the patients underwent a hyperinsulinaemic/hypercarnitinaemic clamp, and had the resting metabolic rate measured before the clamp, 1 hour after the start of insulin infusion, and at the end of the hyperinsulinemic/ hypercarnitinaemic clamp. Vastus lateralis muscle biopsies were taken before and after the clamp (Figure 2).

**Figure 1.**
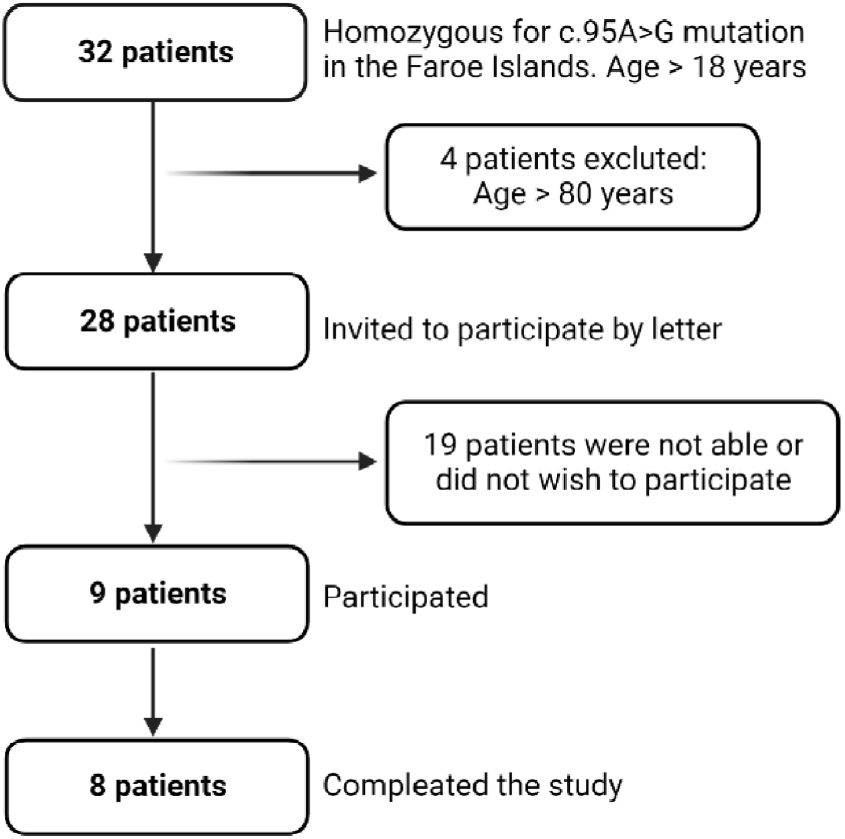
Flowchart of patients with PCD, participating in the study. In the Faroese population, at the time of the study, 28 subjects were known as homozygous for the c.95 A > G mutation and above the age of 18. Nine PCD patients (6 men and 3 women) were included.

**Figure 2.**
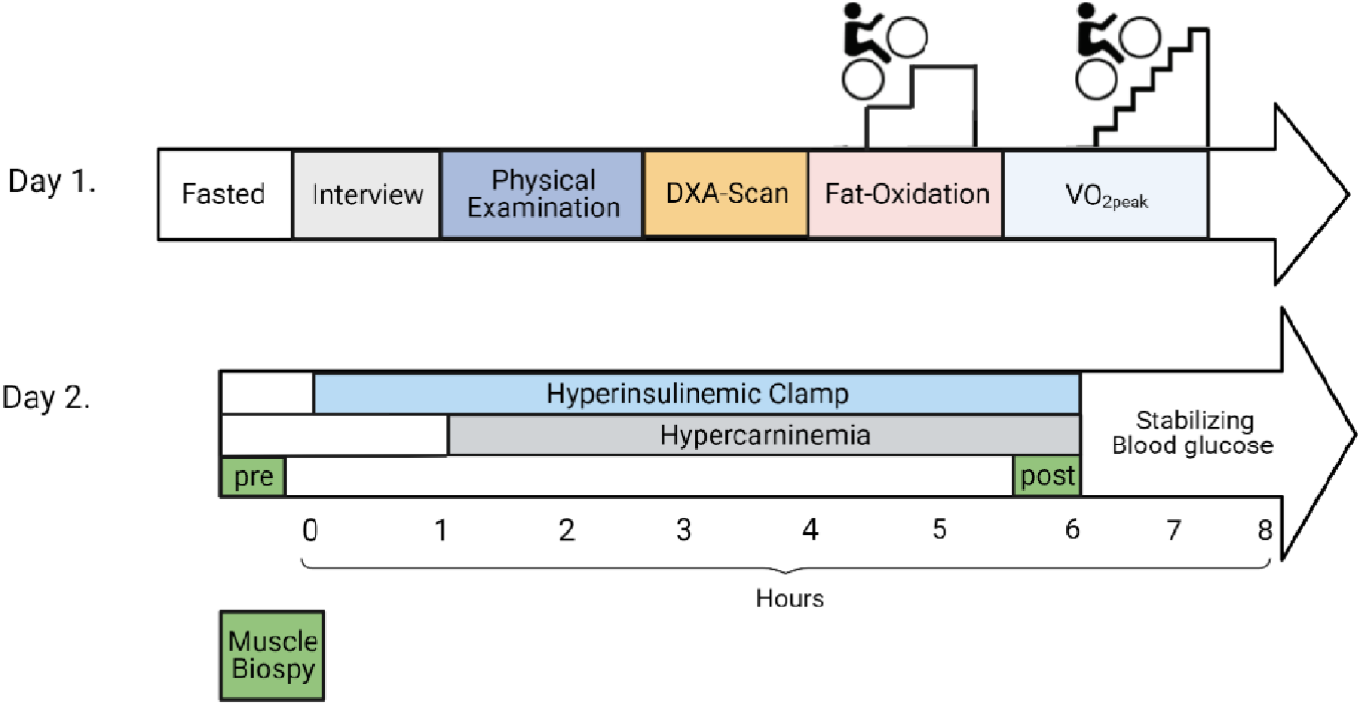
Study design and timeline. Participants reported to the laboratory on two separate occasions, 5-8 days apart. In both cases, the subjects arrived overnight fasted (10 hours). Abbreviations: (DXA) Dual energy X-ray absorptiometry, (VO_2_peak) peak oxygen uptake, (RMR) Resting metabolic rate.

### Experimental procedures

#### Body composition

Each participant had their body composition measured using a dual energy X-ray absorptiometry (DXA) (Norland XR-800, CooperSurgical, Inc., Trumbull, USA). Measurements were obtained while subjects were in the supine position and participants were allowed water in the morning and emptied their bladder before the DXA scan.

#### Exercise test: VO_2_peak and FAT oxidation measurements

To determine FAT oxidation and VO_2_peak, an exercise test was performed on a cycle ergometer (Excalibur Sport, Lode, Groningen, Netherlands) starting out with 1 min rest in a sitting position. This was followed by 10 min at submaximal levels; 4 min at 40/50 watt (w) and 6 min with doubled power output 80/100 w for females and males respectively, using the last 60 s and the equation of Frayn to estimate fat oxidation [30].

Subsequently, the subjects performed a graded protocol in order to determine VO_2_peak, starting at 80/100 w and increasing the power output (20 and 25 w for females and males respectively) every minute until subjects could no longer maintain the desired power output despite verbal encouragement. Oxygen consumption and carbon dioxide production was continuously determined in 15 seconds averages using a mixing chamber analyzing system (model Cosmed, Quark b2, Milan, Italy). Cycling cadence was kept around 80 RPM and a plateau in VO_2_ despite increased power output and a respiratory exchange ratio (RER) >1.15 were used as criteria for VO_2_peak achievement. Gas exchange and ventilation was measured by a sample tube connected to a mixing chamber that was attached to a silicone mask (Hans Rudolph Inc., USA) covering nose and mouth. Prior to every test, the device was calibrated using ambient conditions, the flowmeter was calibrated with a 3-liter syringe and the gas analyzer was calibrated with two gasses of known oxygen and carbon dioxide concentrations. The cycling load at the point of volitional exhaustion (end-power output) was documented as the peak power.

#### Resting metabolic rate

Participants were asked to rest quietly in the supine position and the resting metabolic rate (RMR) was measured by using indirect calorimetry and the ventilated-hood technique (Cosmed, Quark b2, Milan, Italy). RMR was measured before the start of the hyperinsulinemic euglycemic clamp, 40 minutes into the clamp (before starting carnitine infusion) and at the end of the insulin clamp (0). Instruments were calibrated prior to every participant and calibrated against standard mixed reference gases (5 % CO_2_, 16 % O_2_ and the balance N_2_). The measuring time was 20 min and subjects were instructed not to talk and were prevented from falling asleep. RMR was calculated using the formula of de Weir, which is based on measurements of oxygen consumption and carbon dioxide production [31]. Data from each participant has been carefully reviewed based on the notes taken during the experimental measurement to avoid readings during unrest periods.

#### Hyperinsulinemic clamp

All participants were asked to lie down and a cannula was inserted into a superficial vein on their left hand for blood collection [32]. Another cannula was positioned in an antecubital vein in the forearm for insulin and glucose infusion and, furthermore a third cannula was introduced into an antecubital vein in the other arm for L-carnitine infusion.

The subjects underwent a 6 hour euglycaemic hyperinsulinaemic clamp with an insulin (Actrapid, Novo Nordisk, Bagsværd, Denmark) infusion rate of 105mIU/m^2^/min with the aim of achieving a hyperinsulinemic insulin serum level of 150 mIU/l for 6 hours [33]. The blood glucose concentration was maintained by infusion of a 20% glucose solution and after 1h a 5h intravenous infusion of 60 mM L-carnitine solution (MEDICE, Arzneimittel pütter, Iserlohn, Germany) started in parallel with the insulin clamp (0). Carnitine infusion of 15mg/kg was administered for the first 10 minutes in order to quickly increase levels to ∼550 µmol/L followed by 290 minutes of constant infusion at 10 mg/kg/h to maintain hypercarnitinemia. After 6 hours the insulin and carnitine infusions were stopped and the participants had a meal and were kept under observation until their blood glucose levels were stabilized.

#### Blood sampling and analyses

During the insulin clamp, 1 ml of arterialized-venous blood was obtained every 5 minutes for monitoring blood glucose concentration (ABL90 FLEX, plus Radiometer, Denmark). In addition, 2 ml of arterialized-venous blood were obtained every hour for 6 hours, measuring free carnitine, insulin, free fatty acids and glycerol. At baseline and at the end of the insulin clamp an extra 5 ml blood sample was taken where cholesterol (total, LDL and HDL) and C-peptide were measured. After centrifugation, the plasma was removed and stored at -80 °C until further analysis on an automated analyser (Hitachi 912; Roche, Mannheim, Germany) using standard applications. However, Free carnitine concentrations were measured using the radioenzymatic assay described previously by Cederblad et al. [34].

#### Muscle biopsies

All participants had skeletal muscle biopsies taken at two different time-points before and at the end of the insulin clamp. The muscle biopsies were obtained from *m. vastus lateralis* under local anesthesia using the Bergstrøm technique [35, 36]. A piece of the muscle was snap frozen in liquid nitrogen and stored at -80ºC for muscle carnitine content. Another piece was fixed with 2% paraformaldehyde supplemented with 1.2% picric acid for single muscle fibers analysis.

#### Muscle analyses

Free carnitine, acetylcarnitine and long-chain acylcarnitine contents were determined radioenzymatically as described by Cederblad et al [25]. Values were subsequently summed in order to calculate muscle total carnitine.

#### Single muscle fiber isolation and analyses

A fraction of the muscle biopsies were immersed in cold Krebs–Henseleit bicarbonate buffer containing procaine hydrochloride (1g L^-1^) for 5 minutes and, thereafter fixed by immersion in 2% paraformaldehyde supplemented with 1.2% picric acid. Furthermore they were washed in PBS, teased into 10-20 muscle fiber bundles and stored in 50% glycerol/PBS (v/v) for 12 hours at 4 °C before storage at -20 °C, as previously described [37]. Before analysis, muscle fibers were allowed to equilibrate to room temperature and were gently teased apart in PBS into single muscle fibers. The single fibers were used for co-immunostaining of OCTN2 and GLUT4.

#### Co-immunostaining and intracellular analyses of OCTN2 and GLUT4

OCTN2 was labeled using a rabbit polyclonal IgG against α-SLC22A5 (Dil. 1/200, Thermofisher Cat# PA5-102460) and GLUT4 was immunolabeled using a monoclonal mouse IgG antibody (Ab) (Dil. 1/1000, Thermofished Cat# MA1-83191). Fiber-typing was performed by co-immunostaining against α-SERCA2 ATPase, using a monoclonal mouse IgG2a Ab (Dil. 1/400, Thermofisher Cat# MA3-919). Primary antibodies were diluted in immuno buffer (IB: 50 mM glycine, 0.25%, bovine serum albumin, 0.03% saponin and 0.05%, sodium azide in PBS) and, overnight incubated with gentle shacking at room temperature. Single muscle fibers were then washed three times, 30 minutes each, with IB, followed with an overnight incubation with secondary Ab. A goat α-rabbit IgG conjugated to Alexa Flour 568 (Dil. 1/400, Thermofisher, Cat# A11036) was used to immunodetect OCTN2, a goat α-mouse IgG1 conjugated to Alexa Flour 488 (Dil. 1/400, Thermofisher, Cat# A21121) was used to immunodetect GLUT4 and, a goat α-mouse IgG2A conjugated to Alexa Flour 647 (Dil. 1/400, Thermofisher, Cat# A21241). The following day the myonuclei were labelled by incubating the fibers for 5 minutes with DNA specific dye, Hoescht 33342 in IB (Dil. 1/4000, Invitrogen, 33342). After three washes, 30 minutes each, in IB, a final wash with PBS for 10 minutes was performed, and immunostained muscle fibers were mounted in Vectashield mounting medium (Vector Lab, Burlingame, CA, USA, H-1000). Fibers were covered with a coverslip 0.17 ± 0.01 mm thick (#1.5), sealed with nail polish and stored at -20 °C until imaging. Approximately 20-30 fibers were immunostained and analyzed per muscle biopsy.

Image acquisition was performed with a Zeiss LSM700 confocal microscope through a Plan-Apochromat 63x/1.40 objective. Diode lasers were used to excite Hoechst at 405 nm, Alexa Fluor 488 at 488 nm, Alexa Fluor 568 at 555 nm and Alexa Fluor 647 at 639 nm. Pinhole was set to 1 AU for the 488 nm channel and kept for the other three channels to match optical section thickness for all channels. Imaging conditions were set to minimize bleaching, noise and, empty and saturated pixels. In order to investigate whether OCTN2 was present in GLUT4–positive vesicles, GLUT4 and OCTN2 co-localization was assessed in the thin layer of cytoplasm on top of each myonuclei to avoid false co-localization due to random overlap of vesicles in Z. Thus, Z-stacks were collected from the top to the middle of the myonuclei and were used to acquire maximum projections. An illustration of this area is shown in Fiure 3. To avoid several OCTN2 or GLUT4 vesicles stack on top of each other, only the area of cytoplasm between the nuclei and subsarcolemma was chosen for quantification.

Intracellular GLUT4-positive vesicles are known to fuse with the sarcolemma in response to insulin. This results in depletion of the intracellular GLUT4-positive vesicle subsarcolemmal pools [28, 29]. Here, GLUT4 and OCTN2 translocation was quantified by measuring the depletion of both GLUT4-positive and OCTN2-positive vesicles at the subsarcolemmal region over the myonuclei before and after insulin and carnitine clamp. Quantification was performed using CellProfiler [38].

#### Statistical analysis

Data normality was verified by the Kolmogorov-Smirnov test and unpaired *t* test or Welch was used to compare data between groups (0).

A two-way ANOVA (time and group effects) was performed to locate differences in plasma insulin, plasma carnitine, plasma FFA, glycerol, total cholesterol, HDL and LDL, triglycerides and blood glucose concentrations as well as muscle carnitine and long-chain acyl-CoA during the 6 hour euglycaemic hyperinsulinaemic clamp. When a significant main effect was detected, data were further analyzed with student’s paired *t* test using the Šidák correction. To compare data from the resting metabolic rate measurements a mixed model analysis was performed to verify differences between time and groups. For OCTN2 and GLUT4 imaging, a two-way ANOVA was used (post hock test, Šidák’s multiple comparison test).

Statistical significance was considered at *P* < 0.05, and descriptive statistics in text, tables and figures consisted of mean values and mean standard errors (M ± SEM). The analysis were performed with IBM SPSS Statistics 27.0.1.0. and GraphPad™ Prism version 9.0.0 (GraphPad Software Inc) software.

## Results

### Baseline Subjects Characteristics

An overview of subject characteristics are presented in (0). The groups were similar in age, BMI, lean body mass and fat percentage. Furthermore the groups were also similar in plasma FFA, glycerol, total cholesterol, HDL, LDL, triglyceride and plasma free carnitine (0). VO_2_peak was similar between the PCD and control groups, both in terms of absolute values and weight-adjusted. The maximal workload at which VO_2_peak occurred (W_MAX_) was also similar in the two groups (Table 1)

**Table 1.**
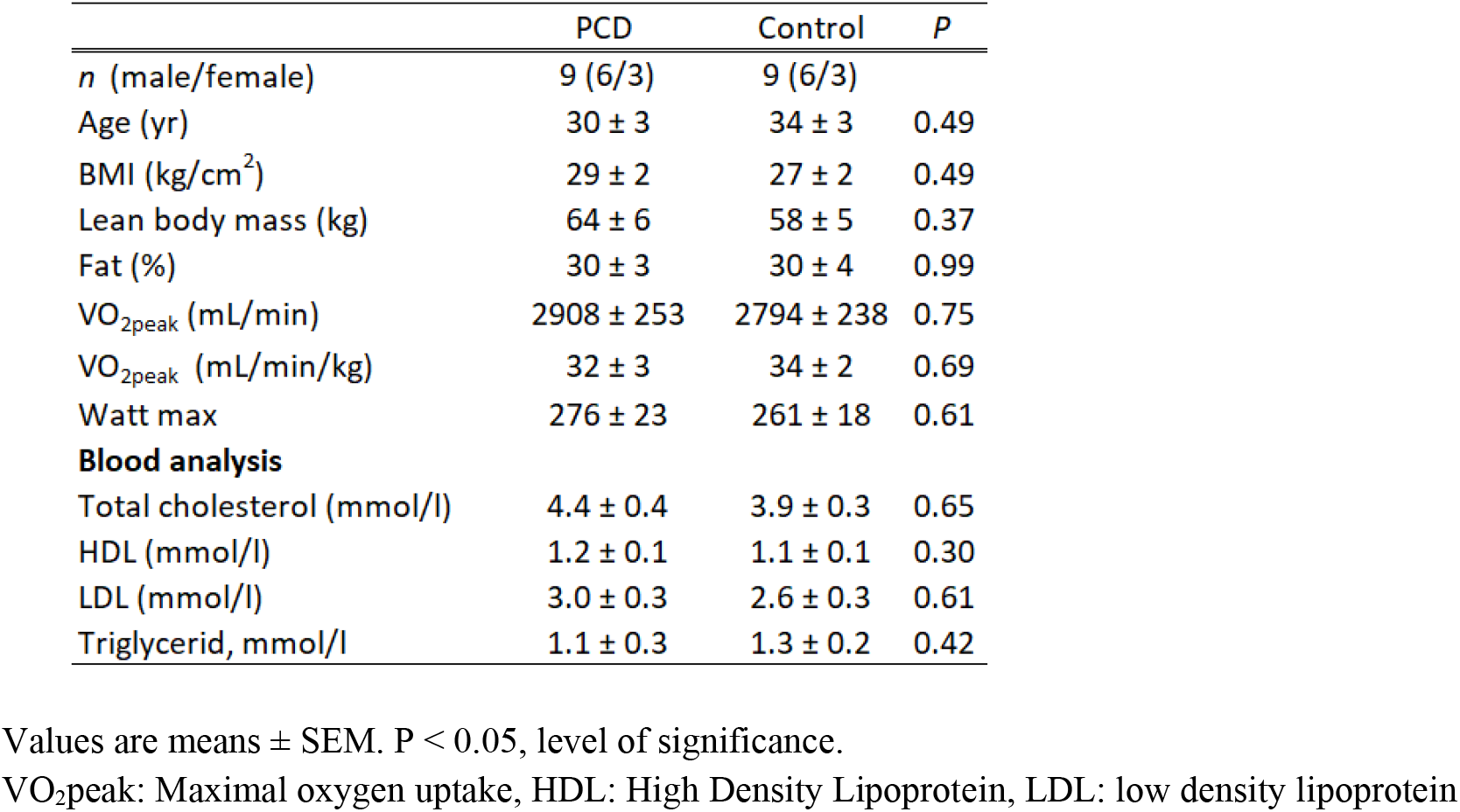
Subject Characteristics

#### Respiration Exchange Ratio (RER) at rest and during exercise

Fat oxidation and RER values from resting fasting state (baseline) and after insulin infusion and the combination of insulin and carnitine infusion, are shown in (0gure 4). At baseline, a significant difference in RER was observed between patients with PCD (0.93 ± 0.03) and controls (RER 0.85 ± 0.01) (*P* = 0.03) but no significant difference was found in fat oxidation rates at rest between patients with PCD (0.04 ± 0.02) and controls (0.06 ± 0.005), (*P* = 0.34). The insulin infusion resulted in an increase of RER in controls (0.96 ± 0.02), reaching levels similar to the once detected in patients with PCD (0.93 ± 0.02) (Figure 4A). RER and the fat oxidation remained stable throughout the rest of the clamp, with no statistically significant difference between groups (*P* > 0.05).

**Figure 3.**
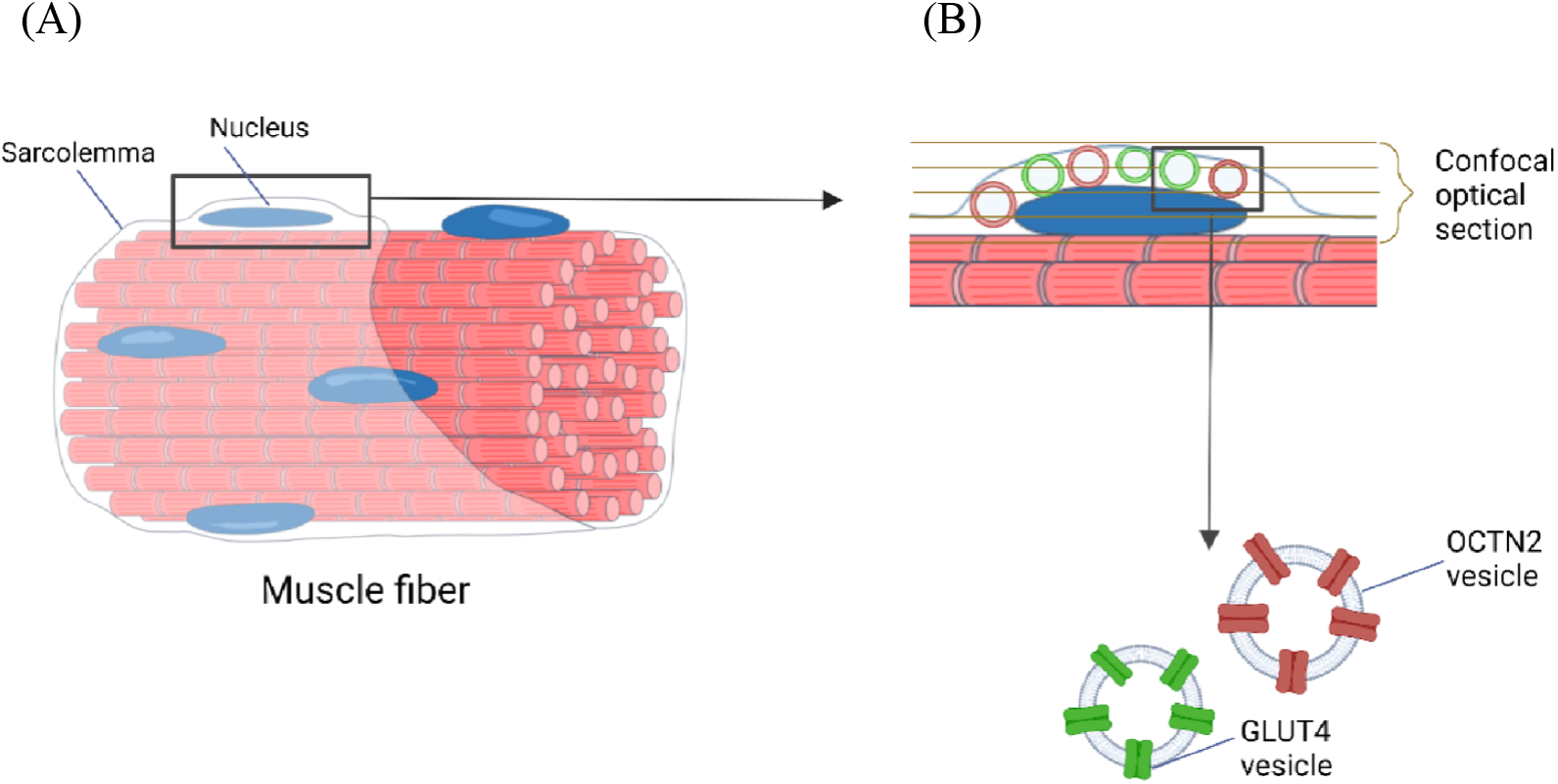
A: Schematic cartoon of single muscle fibers, the sarcolemma, nucleus and B: Intracellular OCTN2 and GLUT4 storage vesicles ready to undergo translocation and fuse with sarcolemma. OCTN2 and GLUT4 positive storage vesicles are visible using confocal microscopic imaging and confocal optical sectioning was performed of the subsarcolemmal region. The depletion storage vesicles was used as an indirect measure of the vesicles translocation to plasma membrane.

**Figure 4.**
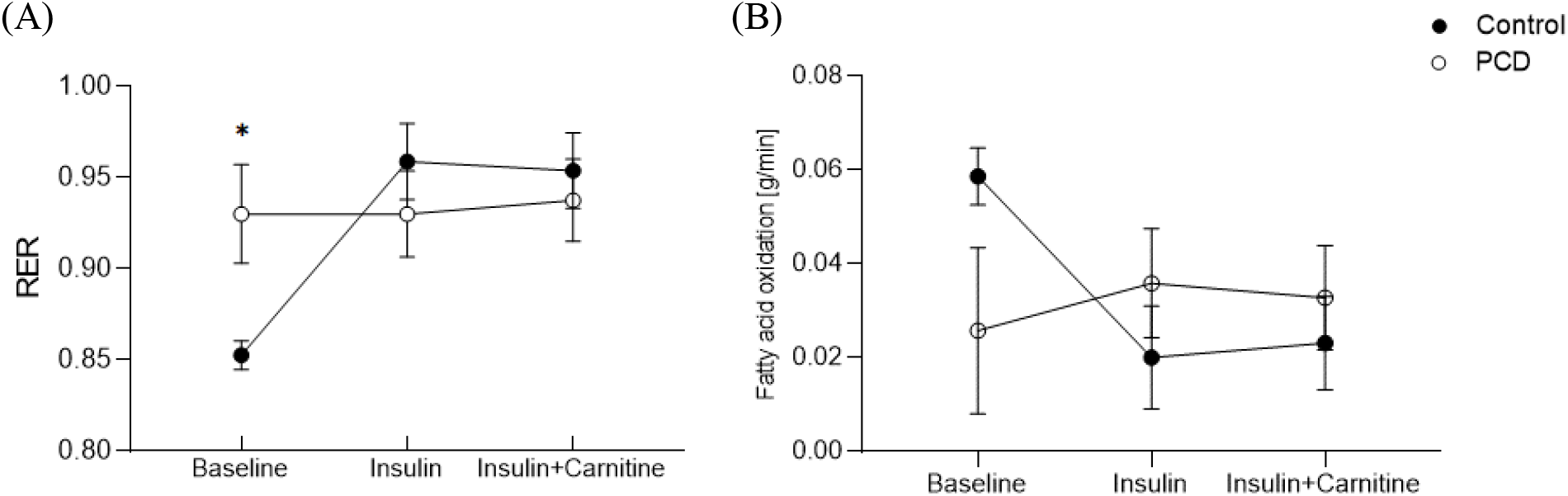
Resting metabolic rate was measured at three time points, at baseline (fasting), after insulin administration (1h) and after insulin (6h) and carnitine (5h) administration combined. Values are mean ± SE and controls (black circles) and patients with PCD (open circles).

No differences were found in RER during exercise at 40/50W (Controls: 0.85 ± 0.04 and PCD: 0.87± 0.06, *P* = 0.84) and at 80/100W (Controls: 0.94 ± 0.04 and PCD: 0.95 ± 0.05, *P* = 0.93) (fig.5A). Furthermore, no differences were found in fat oxidation during exercise at 40/50W (Controls: 0.24 ± 0.05 g/min and PCD: 0.26 ± 0.06 g/min, *P* = 0.95) and at 80/100W (Controls: 0.14 ± 0.04 g/min and PCD: 0.14 ± 0.05 g/min, *P* = 0.99) (0gure 5).

#### Insulin and Carnitine Clamp

The glucose infusion rate (GIR) and the plasma glucose and carnitine concentration during the clamp are shown in 0. At steady state the mean glucose infusion rate was 9.9 ± 1.0 and 9.2 ± 0.8mg/kg/min for controls and patients with PCD respectively (*P* > 0.05). Steady state was defined at time 285-345 min (grey box, Figure 6).

**Figure 5.**
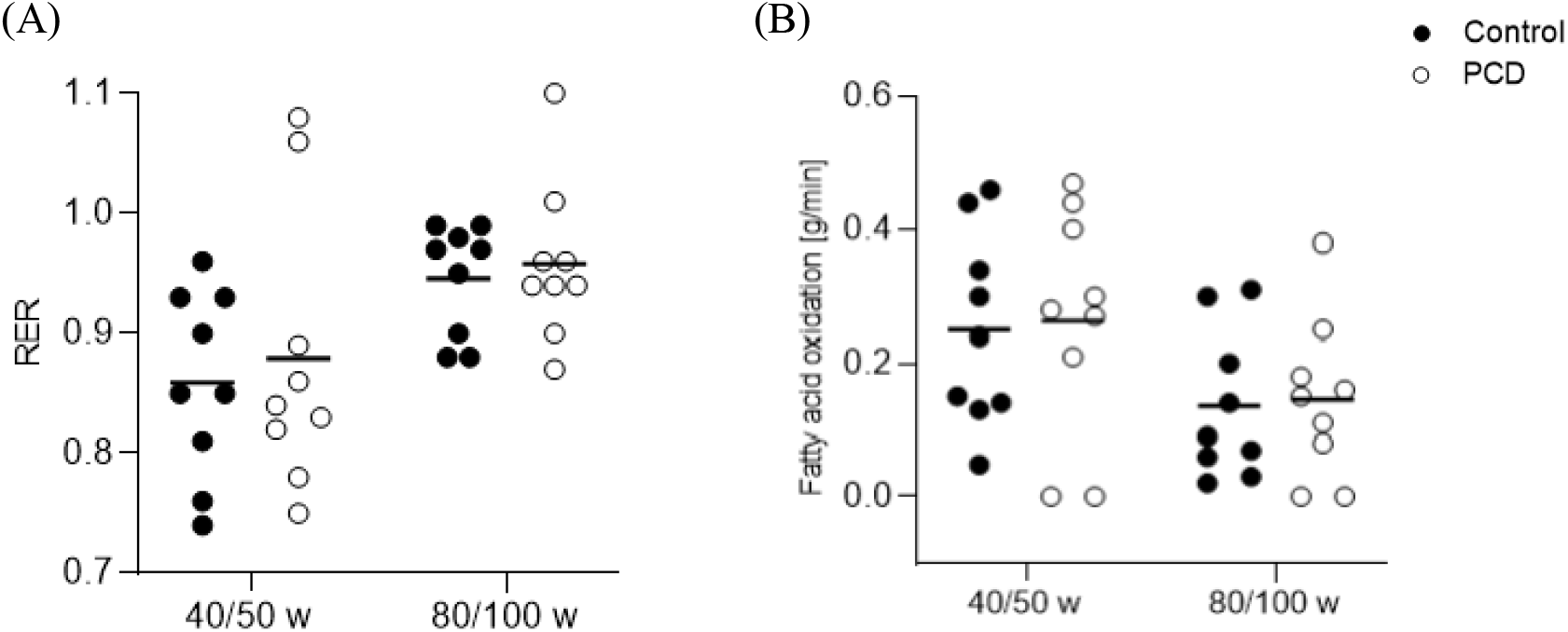
A: RER values and B: fatty acid oxidation rates for controls (black circles) and patients with PCD (open circles) during bike exercise test. All participants worked at two intensities, at 40 and 80 watt for women and 50 and 100 watt for men. Circles indicate individual values and the black line indicates mean values. No significant difference between groups (*P* > 0.05).

**Figure 6.**
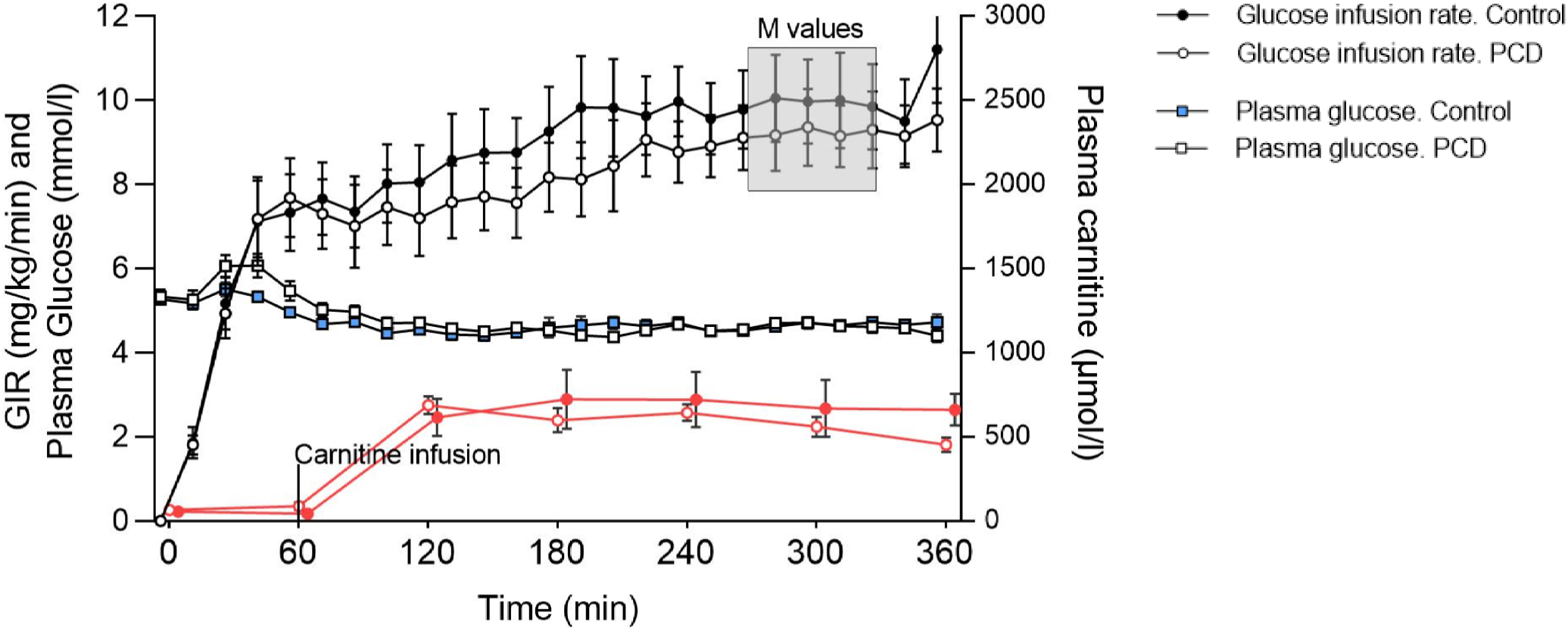
Shows the 6h euglycemic hyperinsulinemic clamps for controls. Left Y-axes shows mean plasma glucose concentrations (mmol/l) and mean glucose infusion rates (mg/kg/min) with the steady state period marked as the 60 minute time period outlined with a rectangle. Right Y-axes indicate carnitine concentrations (µmol/l) during the clamp. Controls are represented as (black, blue and red symbols and patients with PCD as (open symbols). Carnitine was added one hour into the clamp. Values are mean ± SEM and M values were similar between groups (*P* > 0.05).

Table 2 shows biochemical analysis from blood samples collected during the 6 hour euglycemic hyperinsulinaemic clamp. The only significant difference between control subjects and patients with PCD was the free carnitine values at the end of the clamp (*P* < 0.001). Insulin and glucose infusion lead to a clear reduction in both plasma FFA (*P* < 0.01) and glycerol (*P* < 0.01) and a further decrease with the onset of carnitine infusion 60 minutes into the clamp (*P* < 0.01), thus we can see an increase in plasma carnitine and a decrease in glycerol and FFA.

**Table 2.** Substrates in plasma during a 6 hour euglycaemic hyperinsulinaemic clamp

| Time, min | Baseline | 55 | 120 | 180 | 240 | 300 | 360 | *P (time)* |
| --- | --- | --- | --- | --- | --- | --- | --- | --- |
| **Control** |  |  |  |  |  |  |  |  |
| Glucose, mmol/l | 5.4 ± 0.1 | 5.3 ± 0.1 | 4.7 ± 0.1 | 4.9 ± 0.2 | 4.8 ± 0.1 | 4.9 ± 0.1 | 5.0 ± 0.2 | 0.03 |
| Insulin, pmol/l | 86 ± 20 | 1673 ± 108 | 1593 ± 90.5 | 1608 ± 86 | 1556 ± 101 | 1552 ± 105 | 1771 ± 116 | < 0.001 |
| C-Peptide, pmol/l | 876 ± 116 |  |  |  |  |  | 316 ± 55 | < 0.001 |
| FFA, µmol/l | 581 ± 67 | 52 ± 9 | 25 ± 4 | 17 ± 3 | 15 ± 4 | 14 ± 3 | 12 ± 2 | < 0.001 |
| Glycerol, µmol/l | 36.5 ± 11.2 | 5.9 ± 2.1 | 3.4 ± 1.3 | 1.0 ± 0.6 | 1.7 ± 0.9 | 0.6 ± 0.3 | 1.8 ± 0.9 | 0.001 |
| Free Carnitine, µmol/l | 56 ± 10 | 45 ± 8 | 616 ± 110 | 724 ± 175 | 721 ± 164 | 669 ± 169 | 662 ± 95 | < 0.001 |
| **PCD** |  |  |  |  |  |  |  |  |
| Glucose, mmol/l | 5.4 ± 0.2 | 5.9 ± 0.2 | 4.9 ± 0.1 | 4.8 ± 0.1 | 4.9 ± 0.1 | 5.0 ± 0.1 | 4.7 ± 0.1 | < 0.001 |
| Insulin, pmol/l | 90 ± 26 | 1774 ± 140 | 1675 ± 108 | 1672 ± 107 | 1622 ± 116 | 1675 ± 93.0 | 1711 ± 88.3 | < 0.001 |
| C-Peptide, pmol/l | 770 ± 139 |  |  |  |  |  | 368 ± 106 | < 0.001 |
| FFA, µmol/l | 467 ± 62 | 51 ± 6 | 28 ± 6 | 23 ± 5 | 18 ± 4 | 14 ± 3 | 14 ± 4 | 0.005 |
| Glycerol, µmol/l | 23.6 ± 4.2 | 5.1 ± 1.9 | 3.0 ± 0.8 | 2.1 ± 0.8 | 2.4 ± 0.8 | 1.9 ± 0.8 | 2.0 ± 0.7 | 0.002 |
| Free Carnitine, µmol/l | 67 ± 8 | 88 ± 24 | 689 ± 53 | 600 ± 81 | 645 ± 55 | 560 ± 68 | 454 ± 48* | < 0.001 |

Baseline; before the clamp, Time 55 min; insulin infusion, Time 120-360 min; insulin + carnitine infusion, FFA; free fatty acids. Values are means ± SEM, n = 8 PCD patients and 8 controls. *P (time)* is baseline vs. 360 min.
\* $P < 0.0001$ , significantly different from corresponding control group.

#### Muscle carnitine

Due to five hours of hypercarnitinemia and hyperinsulinemia, mean muscle total carnitine tended to increase 1.4 ± 0.6 [19.8±0.6 vs 21.2±0.5 mmol · (kg dry muscle)^-1^], *P* = 0.053 for healthy controls, but remained unchanged for patients with PCD [4.5±0.6 vs 5.3±0.5 · (kg dry muscle)^-1^], *P* = 0.28 (figure 7).

**Figure 7.**
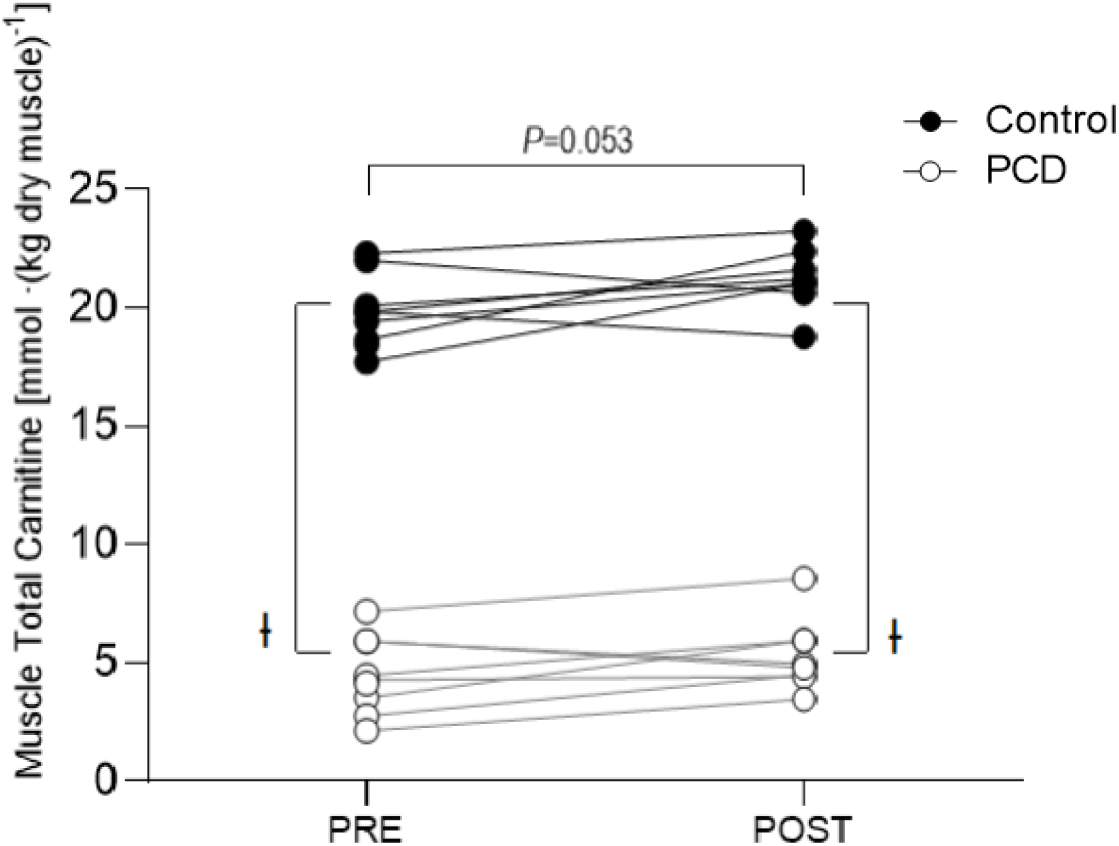
Muscle total carnitine content from skeletal muscle biopsies taken before (PRE) and at the end of the insulin and carnitine infusion (POST). Controls (black circles) and PCD (open circles) and values are from individuals. ł *P* < 0.001, significantly different from control

#### Muscle carnitine profile

Figure 8 visualizes the differences in carnitine moieties (carnitine profile) between controls and patients with PCD, from the biopsies taken before (PRE) and after insulin/carnitine infusion (POST).

**Figure 8.**
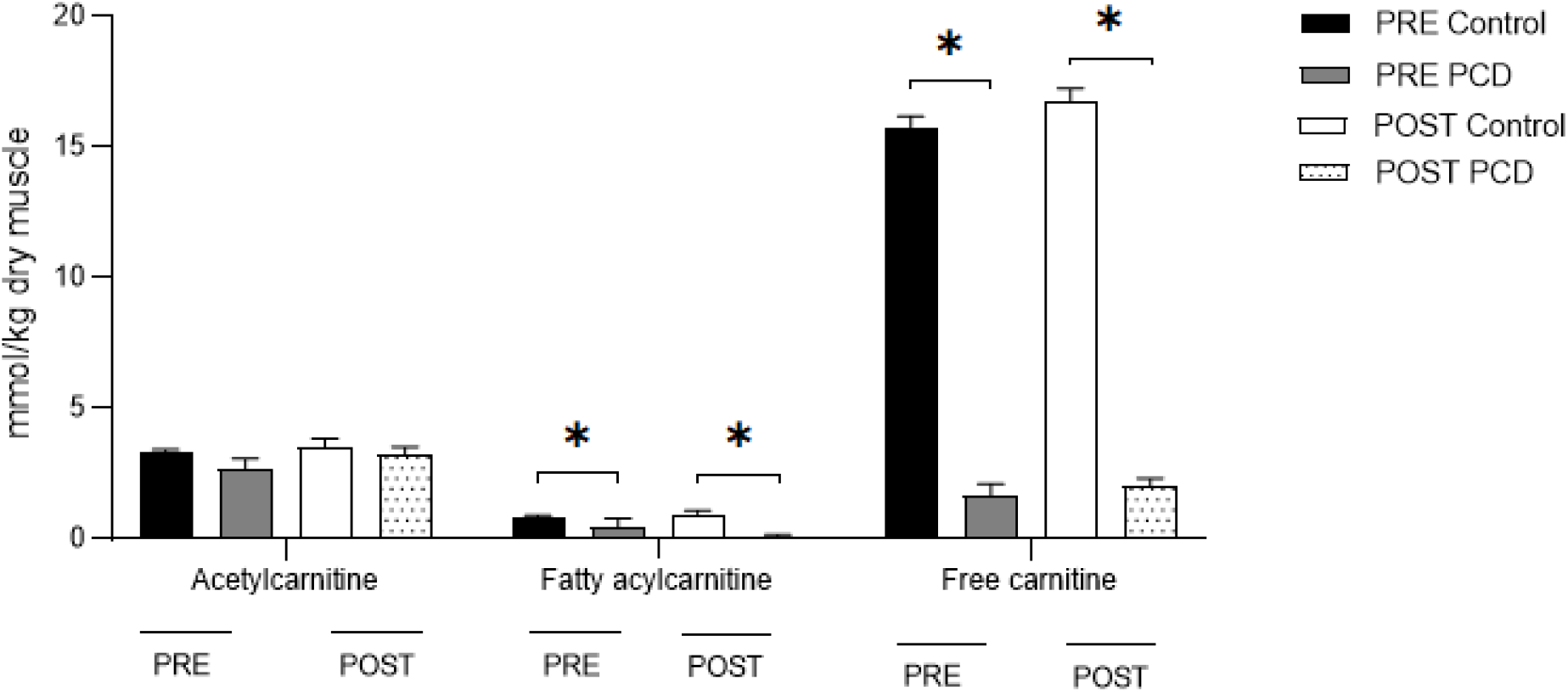
Muscle carnitine profile from biopsies taken before insulin/carnitine infusion (PRE) and after insulin/carnitine infusion (POST). Black bars are Controls PRE, Gray bars are patients with PCD PRE, white bars are controls POST and white/dotted bars are patients with PCD POST. Overall, no significant difference was observed (within groups) due to insulin and carnitine infusion. Patients with PCD had significantly lower muscle free carnitine and fatty acylcarnitne compared to controls, but had normal levels of acetylcarnitine in muscle Values are mean ± SE expressed in mmol/kg dry muscle.* Significant difference between groups, *P* < 0.001.

Overall, insulin and carnitine infusion did not increase muscle carnitine moiety concentrations significantly (*P* > 0.05) (0gure 8). Patients with PCD had significantly lower muscle free carnitine and fatty acylcarnitine compared to controls, both before and after the insulin clamp (*P* < 0.001). However, muscle acetylcarnitine was similar between groups (*P* > 0.05).

#### Insulin-induced OCTN2 and GLUT4 translocation to the sarcolemma

In order to measure insulin-induced GLUT4 and OCTN2 redistribution to the plasma membrane, the depletion of subsarcolemmal vesicles in the thin cytoplasmic volume on top of myonuclei was quantified in type I and type II muscle fibers. In the control group, a significant decrease in OCTN2 vesicles, in the subsarcolemmal area of type I fibers, was observed upon insulin stimulation (*P* = 0.02), indicating that insulin regulates skeletal muscle carnitine uptake through the stimulation of OCTN2 recruitment to the plasma membrane (figure 9 B). There was no decrease in OCTN2 vesicles in type II fibers upon insulin stimulation in the control group (*P* = 0.91, figure 9 C). In contrast, no significant difference in OCTN2 translocation was observed in neither type I fibers (*P* = 0.96) nor type II fibers (*P* = 0.89) from the patients with PCD, indicating that the regulatory mechanism by which insulin upregulates muscle carnitine transport might be impaired in patients with PCD (Figure 9 B, C).

**Figure 9.**
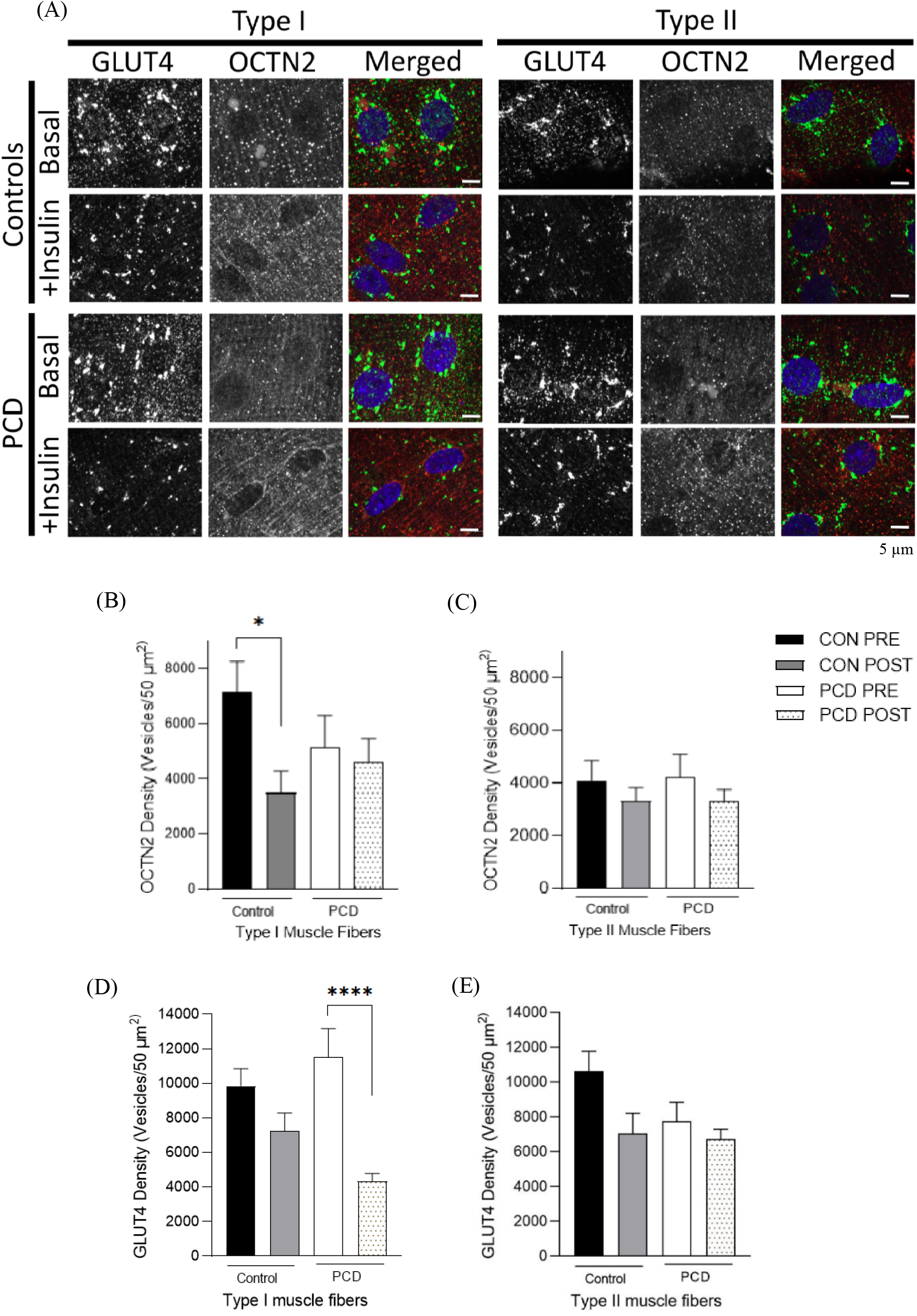
(A) Single muscle fiber co-stained against GLUT4 (green), OCTN2 (red) and nuclei (Blue) at basal and after insulin and carnitine stimulation. (B) Mean OCTN2 values are plotted as PRE control (black), POST insulin stimulated control (gray), PCD PRE (white) and POST insulin stimulated PCD (White with pattern) for type I muscle fibers and (C) type II muscle fibers. (D and E) show mean GLUT4 density in type I fibers and type II fiber in healthy controls and patients with PCD, PRE and POST insulin and carnitine stimulation. Values are mean ± SE expressed in OCTN2 density/50µm^2^. * *P* < 0.05, **** *P* < 0.0001.

There was no significant depletion of GLUT4-positive vesicles in the subsarcolemmal area in type I fibers from control subjects. A tendency to a decrease in GLUT4 positive vesicles was observed upon insulin stimulation in type II fibers (*P* = 0.07) from control subjects, while a significant depletion of GLUT4 positive vesicles was detected in PCD type I muscle fibers (*P* < 0.0001). PCD type II muscle fibers showed no depletion of GLUT4-positive vesicles in the subsarcolemmal area (Figure 9 D, E).

## Discussion

The first question, this study sought to investigate was whether the combination of hypercarnitinaemia and hyperinsulinaemia could increase skeletal muscle carnitine uptake in patients with PCD. In this respect, we found no significant increase in patients with PCD (*P* = 0.28) and a tendency for an increase in controls (*P* = 0.053). The lack of significant increase of insulin induced carnitine uptake in PCD is consistent with the impairment in insulin induced translocation of OCTN2 transporters from intracellular storage to the sarcolemma. However, it could be interesting to see if the muscle carnitine content would increase in PCD with increased insulin combined with carnitine supplementation over a longer period of time as demonstrated by Wall et al. in healthy participants [39]. This study showed that L-carnitine in combination with a beverage containing carbohydrate (in order to stimulate insulin-mediated muscle carnitine accumulation) for up to 24 weeks, increased skeletal muscle total carnitine content by around 20% in young healthy men.

The muscle biopsies also showed that the muscle carnitine profile was different between groups. With patients with PCD having lower free carnitine and acylcarnitine, but the acetylcarnitine levels were normal and representing around 60% of total carnitine for PCD and around 16% for controls. This observation holds significance as it implies that the role of free carnitine to act as an acetyl group buffer remains intact in patients with PCD. Previous research has demonstrated a reduction in acetylcarnitine levels during clamp procedures [25], suggesting its association with fat oxidation - a process inhibited by insulin. Surprisingly, this pattern did not align with the findings of our study, including those from the control group, presenting a notable discrepancy that warrants further investigation.

Another aim of this study was to investigate whether insulin-induced upregulation of carnitine in skeletal muscle is dependent on sarcolemmal recruitment of OCTN2. In this regard, the results show a significant decrease in OCTN2 vesicles, in the subsarcolemmal area, observed upon insulin stimulation in type I fibers (*P* = 0.02) of the control group, indicating that insulin regulates skeletal muscle carnitine uptake through the stimulation of OCTN2 recruitment to the plasma membrane (Figure 9). However, this decrease in OCTN2 vesicles, in the subsarcolemmal area, due to insulin stimulation, was not observed in patients with PCD (homozygous for the p.N32S mutation), indicating that their regulatory mechanism might be impaired (*P* < 0.05). These results reflect those of Filippo et al. where they identify impaired maturation of OCTN2 transporters to plasma membrane in patients with PCD, homozygote for the mutations P46S and R83L associated with the OCTN2 carnitine transporter [10, 40]. However, the P46S and R83L mutations are located in the extracellular loop between the first two transmembrane domains of the transporter and the p.N32S mutation (investigated in present study) is located in the first transmembrane domain [10, 41, 42]. While earlier research has identified two mutations within the first transmembrane domain in PCD patients, leading to low carnitine levels in the body, the precise role of this domain has remained elusive. As far as our knowledge extends, this is the first study to indicate that the first transmembrane domain could play a role in translocation of OCTN2 to the plasma membrane in healthy controls, contributing to novel insights to the field. Notably, an impairment in translocation could result in fever OCTN2 transporters in sarcolemma, which could limit the uptake of carnitine into muscle and limits the transport of LCFA across the inner mitochondrial membrane, affecting the rate of β-oxidation and affecting the cellular metabolism and energy production. However, Stanley and colleagues hypothesized that if the intracellular concentration of carnitine is greater than 4% of normal, fatty acid oxidation in muscle is not compromised [21].

In present study we do not investigate the function of the transporter per se, but the translocation from transmembrane storage pools to the membrane. Previous study by Filippo et al., has shown other mutations associated with the OCTN2 transporter, to impair transporter function without impairing translocation to the plasma membrane [43]. Furthermore others have measured the carnitine uptake in fibroblast cells to be impaired in PCD, without assessment of the amount of OCTN2 transporters in the cell membrane [44]. Therefore, it is yet to be investigated whether PCD (homozygous for the p.N32S mutation) is caused by fewer transporters in the membrane alone or from both fewer and defective transporters. Regardless of a defect in insulin induced translocation of OCTN2 positive vesicles in PCD skeletal muscle there are other mechanisms that could stimulate L-carnitine uptake. Animal studies by Furuicihi et al. [45] have shown that skeletal muscle contraction increases L-carnitine uptake via translocation of OCTN2 to sarcolemma and other animal studies do also support the hypothesis that exercise may activate transmembrane influx of carnitine [46, 47]. Thus, it is a possibility that muscle contraction could induce translocation of OCTN2 to sarcolemma in PCD skeletal muscle, which is supported by the fact that we found no difference in fatty acid oxidation, between groups, during short-term exercise.

Present study did also set out to measure insulin-induced recruitment of GLUT4 to sarcolemma (0). This was done by measuring the depletion of GLUT4 positive vesicles from intracellular storage, upon insulin stimulation and the results showed a tendency to a decrease in type II fibers of controls (*P* = 0.07) and a significant depletion in type I fibers of PCD (*P* < 0.001). Indicating that insulin stimulates GLUT4 translocation to sarcolemma in PCD type I fibers. Surprisingly, insulin stimulation did not induce a significant translocation of GLUT4 in control fibers, (however a tendency was found in type II fibers). This is surprising as it is well established in the literature that insulin stimulates cellular glucose uptake via the GLUT4 transporters [28, 29, 48-50].

The results from GIR (>5 mg/kg/min), measured during the hyperinsulinemic clamp support the fact that none of the groups showed signs of insulin resistance, with no significant difference between PCD and control group (*P* = 0.63). Confocal microscopy (0) show that the OCTN2 and GLUT4 transporters have different localization in skeletal muscle, which could indicate that they are in different transport vesicles. Furthermore, the translocation quantification showing that patients with PCD have an impaired insulin stimulated OCTN2 translocation, but no impairment in the insulin stimulated GLUT4 translocation, also supports that they are located in different transport vesicles. Interestingly, the p.N32S mutation in the OCTN2 is in a domain that is highly conserved between OCTN2 and GLUT4 and is known to be involved in insulin response [49].

Another finding of the present study was the resting metabolic rate where we found the RER values for PCD to be significantly higher than controls, indicating that patients with PCD were more dependent on carbohydrate metabolism than controls at resting fasting state (basal) (0gure 4A). This observation can be attributed to the necessity of free carnitine in facilitating the transport of long-chain fatty acids into mitochondria, which is essential for fatty acid oxidation. However when considering the absolute values with the formula of Frayn et al. [30], there was no significant difference in fat oxidation (g/min) between groups (Figure 4B), however tendency (*P* = 0.95) was found. Interestingly, acetylcarnitine concentrations in muscle from PCD were found to be within normal range suggesting that carnitine effectively serves as an acetyl group buffer.

As far as we know, only one similar study has been conducted which found no differences in fatty acid oxidation between patients with PCD and healthy controls at rest [22]. However, considering that the measurements are whole-body fatty acid oxidation, one plausible explanation is that the impaired fat oxidation may not be occurring in skeletal muscle.

The difference in RER found at basal, disappears at the start of the insulin clamp as the control RER values increase by 13% and reach the same levels as the patients with PCD and both groups are predominantly metabolizing carbohydrates. The increase could be expected because blood glucose levels were kept stable during the clamp by infusing a 20% glucose solution.

Interestingly, our investigations found no difference in fatty acid oxidation between groups, during 10 min submaximal exercise; (4 min at 40/50 w and 6 min 80/100 w) (0) and high intensity exercise (VO2max). Notably, the ability of patients with PCD, who exhibit low muscle carnitine levels, to perform perform high intensity exercise is interesting as carnitine has been shown to play a role as an acetyl buffer during high intensity exercise. Supporting this we found that the muscle acetylcarnitine levels of the PCD patients were comparable to the controls, indicating that the buffering role of carnitine is intact. The role of carnitine during high intensity exercise has been explored in previous research. For instance in a study by Sahlin et al, it was noted that muscle free carnitine content, which was approximately 16 mmol (kg dry muscle)^-1^ at rest, decreased to about one-third of the initial value after a 15 min exercise at 75% and 100% VO2max in healthy volunteers. This reduction was primarily attributed to the formation of acetylcarnitine [51]. Additionally, in a study by Bouitbir et al limited physical performance was observed in carnitine depleted rats performing high intensity exercise [52]. Therefore, given that free carnitine levels tend to decrease during high-intensity exercise, one might expect individuals with initially low free carnitine levels, such as PCD patients, to have restricted capabilities for high-intensity exercise. However, our present study demonstrates that high-intensity exercise is achievable even with very low muscle carnitine concentrations, as PCD patients. This is further supported by Madsen et al, who also found no significant difference between PCD patients and the control group when performing high-intensity exercise (VO2max). It should be noted that the patients in the present study were receiving L-carnitine supplementation, a factor shared with the mentioned study by Madsen et al.

In summary, we found that the combination of hypercarnitinaemia with hyperinsulinemia did not increase skeletal muscle total carnitine levels significantly in patients with PCD. Furthermore we found that the skeletal muscle profile is different in patients with PCD, with low free and long-chain acylcarnitine and normal levels of acetylcarnitine, compared to healthy controls. Confocal microscopy indicated that insulin stimulates OCTN2 recruitment to the plasma membrane in type I fibers in healthy controls, a mechanism that seems to be impaired in patients with PCD, however the results indicated that insulin stimulates GLUT4 recruitment to the plasma membrane in type I fiber in PCD and a tendency for recruitment of GLUT4 in type II fibers in controls. The hyperinsulinemic clamp showed that none of the groups suffered from insulin resistance. Present study does also hypothesize that OCTN2 and GLUT4 transporters are located in different transport vesicles, since they react differently to insulin stimulation and results from indirect calorimetry indicate that patients with PCD metabolize energy reserves differently and depend more on carbohydrate oxidation at rest. However no difference was found between PCD and control group during short-term exercise and patients with PCD are able to perform high intensity exercise.

### Limitations

A limitation of this study was the fact that the patients with PCD had taken their L-carnitine supplementation as normal, including the morning of the insulin and L-carnitine infusion. This could have an influence on the muscle carnitine levels at basal. An alternative could have been a pause in the L-carnitine supplements, but this would have included a greater risk for the patients and require monitoring of the patients in a hospital setting.

## Acknowledgements

The project was supported by funds from the Faroese Research Consul, the AP Møller Foundation, The Faroese CTD association (CTD Felagið) and the Center for Healthy Aging, University of Copenhagen.

We thank the individuals with PCD and the controls who volunteered to participate in this study. We would also like to thank Hildegunn S. Thomsen, Halla Weihe Reinert and Elsa Splidt Jacobsen, for their help during the experiments and Debes Christiansen (the Faroese Food and Veterinary Authority) for genetic analysis. Furthermore we would like to thank Regitze Kraunsøe and Jeppe Bach for technical assistance, the National Hospital of the Faroe Island for supporting the project and Ann E. Østerø for providing us with space and assistance at their clinical laboratory.

## Notes

### Competing Interest Statement

The authors have declared no competing interest.

## References

1. Joensen, F., E.U. Steuerwald, and N.H. Rasmussen, [Three congenital metabolic diseases in the Faeroe Islands. Incidence, clinical and molecular genetic characteristics of Faeroese children with glycogen storage disease type IIIA, carnitine transporter deficiency and holocarboxylase synthetase deficiency]. Ugeskr Laeger, 2006. 168(7): p. 667–70.

2. Osá, E. and H. Simonsen, [Carnitine transporter deficiency in two Faeroese children]. Ugeskr Laeger, 2004. 166(50): p. 4612–3.

3. Rasmussen, J., et al., Increased risk of sudden death in untreated primary carnitine deficiency. J Inherit Metab Dis, 2020. 43(2): p. 290–296.

4. Rasmussen, J., et al., Carnitine levels in 26,462 individuals from the nationwide screening program for primary carnitine deficiency in the Faroe Islands. J Inherit Metab Dis, 2014. 37(2): p. 215–22.

5. Rasmussen, J., et al., Primary carnitine deficiency and pivalic acid exposure causing encephalopathy and fatal cardiac events. J Inherit Metab Dis, 2013. 36(1): p. 35–41.

6. Lund, A.M., et al., Carnitine transporter and holocarboxylase synthetase deficiencies in The Faroe Islands. J Inherit Metab Dis, 2007. 30(3): p. 341–9.

7. De Biase, I., et al., Primary Carnitine Deficiency Presents Atypically with Long QT Syndrome: A Case Report. JIMD Rep, 2012. 2: p. 87–90.

8. El-Hattab, A.W., et al., Maternal systemic primary carnitine deficiency uncovered by newborn screening: clinical, biochemical, and molecular aspects. Genet Med, 2010. 12(1): p. 19–24.

9. Dambrova, M., et al., Acylcarnitines: Nomenclature, Biomarkers, Therapeutic Potential, Drug Targets, and Clinical Trials. Pharmacol Rev, 2022. 74(3): p. 506–551.

10. Longo, N., M. Frigeni, and M. Pasquali, Carnitine transport and fatty acid oxidation. Biochim Biophys Acta, 2016. 1863(10): p. 2422–35.

11. Longo, N., C. Amat di San Filippo, and M. Pasquali, Disorders of carnitine transport and the carnitine cycle. Am J Med Genet C Semin Med Genet, 2006. 142c(2): p. 77–85.

12. Rebouche, C.J., Kinetics, pharmacokinetics, and regulation of L-carnitine and acetyl-L-carnitine metabolism. Ann N Y Acad Sci, 2004. 1033: p. 30–41.

13. Steiber, A., J. Kerner, and C.L. Hoppel, Carnitine: a nutritional, biosynthetic, and functional perspective. Mol Aspects Med, 2004. 25(5-6): p. 455–73.

14. Adeva-Andany, M.M., et al., Significance of l-carnitine for human health. IUBMB Life, 2017. 69(8): p. 578–594.

15. Gnoni, A., et al., Carnitine in Human Muscle Bioenergetics: Can Carnitine Supplementation Improve Physical Exercise? Molecules, 2020. 25(1).

16. Pettegrew, J.W., J. Levine, and R.J. McClure, Acetyl-L-carnitine physical-chemical, metabolic, and therapeutic properties: relevance for its mode of action in Alzheimer’s disease and geriatric depression. Mol Psychiatry, 2000. 5(6): p. 616–32.

17. Schulz, H., Regulation of fatty acid oxidation in heart. J Nutr, 1994. 124(2): p. 165–71.

18. Christensen, E. and J. Vikre-Jørgensen, Six years’ experience with carnitine supplementation in a patient with an inherited defective carnitine transport system. J Inherit Metab Dis, 1995. 18(2): p. 233–6.

19. Yasuro Furuichi, N.G.-I.a.N.L.F., Role of carnitine acetylation in skeletal muscle. J Phys Fitness Sports Med, 3(2), 2014: p. 163–168.

20. Lamhonwah, A.M., et al., Novel OCTN2 mutations: no genotype-phenotype correlations: early carnitine therapy prevents cardiomyopathy. Am J Med Genet, 2002. 111(3): p. 271–84.

21. Stanley, C.A., et al., Chronic cardiomyopathy and weakness or acute coma in children with a defect in carnitine uptake. Ann Neurol, 1991. 30(5): p. 709–16.

22. Madsen, K.L., et al., L-Carnitine Improves Skeletal Muscle Fat Oxidation in Primary Carnitine Deficiency. J Clin Endocrinol Metab, 2018. 103(12): p. 4580–4588.

23. Rasmussen, J., et al., Carnitine levels in skeletal muscle, blood, and urine in patients with primary carnitine deficiency during intermission of L-carnitine supplementation. JIMD Rep, 2015. 20: p. 103–11.

24. Stephens, F.B., et al., A threshold exists for the stimulatory effect of insulin on plasma L-carnitine clearance in humans. Am J Physiol Endocrinol Metab, 2007. 292(2): p. E637-41.

25. Stephens, F.B., et al., Insulin stimulates L-carnitine accumulation in human skeletal muscle. Faseb j, 2006. 20(2): p. 377–9.

26. Piper, R.C., et al., GLUT-4 NH2 terminus contains a phenylalanine-based targeting motif that regulates intracellular sequestration. J Cell Biol, 1993. 121(6): p. 1221–32.

27. Araki, S., et al., Subcellular trafficking kinetics of GLU4 mutated at the N- and C-terminal. Biochem J, 1996. 315 ( Pt 1)(Pt 1): p. 153–9.

28. Lauritzen, H.P. and J.D. Schertzer, Measuring GLUT4 translocation in mature muscle fibers. Am J Physiol Endocrinol Metab, 2010. 299(2): p. E169-79.

29. Lauritzen, H.P., Insulin- and contraction-induced glucose transporter 4 traffic in muscle: insights from a novel imaging approach. Exerc Sport Sci Rev, 2013. 41(2): p. 77–86.

30. Frayn, K.N., Calculation of substrate oxidation rates in vivo from gaseous exchange. J Appl Physiol Respir Environ Exerc Physiol, 1983. 55(2): p. 628–34.

31. Weir, J.B., New methods for calculating metabolic rate with special reference to protein metabolism. J Physiol, 1949. 109(1-2): p. 1–9.

32. Gallen, I.W. and I.A. Macdonald, Effect of two methods of hand heating on body temperature, forearm blood flow, and deep venous oxygen saturation. Am J Physiol, 1990. 259(5 Pt 1): p. E639-43.

33. DeFronzo, R.A., J.D. Tobin, and R. Andres, Glucose clamp technique: a method for quantifying insulin secretion and resistance. Am J Physiol, 1979. 237(3): p. E214–23.

34. Cederblad, G., O. Finnstrom, and J. Martensson, Urinary excretion of carnitine and its derivatives in newborns. Biochem Med, 1982. 27(2): p. 260–5.

35. Bergstrom, J., Percutaneous needle biopsy of skeletal muscle in physiological and clinical research. Scand J Clin Lab Invest, 1975. 35(7): p. 609–16.

36. Bergstrom, J., Muscle-biopsy needles. Lancet, 1979. 1(8108): p. 153.

37. Dahl, R., et al., Three-dimensional reconstruction of the human skeletal muscle mitochondrial network as a tool to assess mitochondrial content and structural organization. Acta Physiol (Oxf), 2015. 213(1): p. 145–55.

38. Carpenter, A.E., et al., CellProfiler: image analysis software for identifying and quantifying cell phenotypes. Genome Biol, 2006. 7(10): p. R100.

39. Wall, B.T., et al., Chronic oral ingestion of L-carnitine and carbohydrate increases muscle carnitine content and alters muscle fuel metabolism during exercise in humans. J Physiol, 2011. 589(Pt 4): p. 963–73.

40. Filippo, C.A., O. Ardon, and N. Longo, Glycosylation of the OCTN2 carnitine transporter: study of natural mutations identified in patients with primary carnitine deficiency. Biochim Biophys Acta, 2011. 1812(3): p. 312–20.

41. Li, F.Y., et al., Molecular spectrum of SLC22A5 (OCTN2) gene mutations detected in 143 subjects evaluated for systemic carnitine deficiency. Hum Mutat, 2010. 31(8): p. E1632-51.

42. Amat di San Filippo, C., Y. Wang, and N. Longo, Functional domains in the carnitine transporter OCTN2, defective in primary carnitine deficiency. J Biol Chem, 2003. 278(48): p. 47776–84.

43. Amat di San Filippo, C. and N. Longo, Tyrosine residues affecting sodium stimulation of carnitine transport in the OCTN2 carnitine/organic cation transporter. J Biol Chem, 2004. 279(8): p. 7247–53.

44. Rasmussen, J., et al., Residual OCTN2 transporter activity, carnitine levels and symptoms correlate in patients with primary carnitine deficiency. Mol Genet Metab Rep, 2014. 1: p. 241–248.

45. Furuichi, Y., et al., Muscle contraction increases carnitine uptake via translocation of OCTN2. Biochem Biophys Res Commun, 2012. 418(4): p. 774–9.

46. Vukovich, M.D., D.L. Costill, and W.J. Fink, Carnitine supplementation: effect on muscle carnitine and glycogen content during exercise. Med Sci Sports Exerc, 1994. 26(9): p. 1122–9.

47. Broderick, T.L., et al., Biosynthesis of the Essential Fatty Acid Oxidation Cofactor Carnitine Is Stimulated in Heart and Liver after a Single Bout of Exercise in Mice. J Nutr Metab, 2018. 2018: p. 2785090.

48. Klip, A., T.E. McGraw, and D.E. James, Thirty sweet years of GLUT4. J Biol Chem, 2019. 294(30): p. 11369–11381.

49. Huang, S. and M.P. Czech, The GLUT4 glucose transporter. Cell Metab, 2007. 5(4): p. 237–52.

50. Jaldin-Fincati, J.R., et al., Update on GLUT4 Vesicle Traffic: A Cornerstone of Insulin Action. Trends Endocrinol Metab, 2017. 28(8): p. 597–611.

51. Sahlin, K., Muscle carnitine metabolism during incremental dynamic exercise in humans. Acta Physiol Scand, 1990. 138(3): p. 259–62.

52. Bouitbir, J., et al., Impaired Exercise Performance and Skeletal Muscle Mitochondrial Function in Rats with Secondary Carnitine Deficiency. Front Physiol, 2016. 7: p. 345.

